# Burst activation of dopamine neurons produces prolonged post-burst availability of actively released dopamine

**DOI:** 10.1101/125062

**Authors:** Sweyta Lohani, Adria K. Martig, Suzanne M. Underhill, Alicia DeFrancesco, Melanie J. Roberts, Linda Rinaman, Susan Amara, Bita Moghaddam

## Abstract

Both phasic and tonic modes of neurotransmission are implicated in critical functions assigned to dopamine. In learning, for example, sub-second phasic responses of ventral tegmental area (VTA) dopamine neurons to salient events serve as teaching signals, but learning is also interrupted by dopamine antagonists administered minutes after training. Our findings bridge the multiple timescales of dopamine neurotransmission by demonstrating that burst stimulation of VTA dopamine neurons produces a prolonged post-burst increase (> 20 min) of extracellular dopamine in nucleus accumbens and prefrontal cortex. This elevation is not due to spillover from the stimulation surge but depends on impulse flow-mediated dopamine release. We identified Rho-mediated internalization of dopamine transporter as a mechanism responsible for prolonged availability of actively released dopamine. These results demonstrate that phasic and tonic dopamine neurotransmission can be a continuum and may explain why both modes of signaling are critical for motivational and cognitive functions associated with dopamine.

## Introduction

Dopamine neurons project extensively to striatal and cortical regions where they mediate a diverse array of functions, including reward-related learning, drug-related synaptic plasticity, working memory and motivation ^1-6^. Dopamine neurons respond with brief and phasic bursts of action potentials to salient events, such as reward-predictive stimuli and unexpected reward delivery ^7,8^, and these bursts cause phasic dopamine release in terminal regions, including the prefrontal cortex (PFC) and the nucleus accumbens (NAc) ^9-12^. Influential theories suggest these phasic responses serve as teaching signals in reinforcement learning ^7,13,14^ and/or motivational signals that mediate cue-triggered seeking of rewards ^1,4^.

While fast-scan cyclic voltammetry (FSCV) studies have characterized rapid and transient increases in extracellular dopamine concentration, consistent with phasic bursting of dopamine neurons, in response to salient events in a behavioral task ^15-18^, microdialysis studies have demonstrated sustained increases in cortical and striatal extracellular dopamine that persist for many minutes (even up to an hour) after the completion of appetitive learning, instrumental behavior and other cognitive tasks ^19-23^. The prolonged post-session dopamine increase occurs despite the fact that animals are no longer receiving rewards and/or being presented with salient stimuli. Importantly, dopamine antagonists administered either systemically or directly into PFC and striatum minutes after behavioral training can disrupt memory consolidation, demonstrating the necessity of post-training dopamine signaling ^24-26^. This slower mode of dopamine signaling has been attributed to the so-called “tonic” dopamine release, which is hypothesized to be mediated by mechanisms that are entirely distinct from those that drive phasic activity of dopamine neurons and release ^27-29^. But given that learning and most behavioral functions assigned to dopamine depend on both fast and slow modes of neurotransmission, we hypothesized that these phasic and tonic dopamine states might instead comprise a continuum of the same activation process in an awake and active animal. In that case, phasic activation of dopamine neurons could trigger secondary events that enhance the slow (tonic) activation of dopamine levels. Thus, we designed experiments to determine whether bursting of dopamine neurons results in post-burst sustained elevation of extracellular levels of dopamine in terminal regions and, if so, what mechanisms sustain this elevation.

Dopamine’s effects on reward-related learning, motivation and cognitive constructs such as attention and working memory are primarily mediated by dopamine neurons in ventral tegmental area (VTA) and their projections to the NAc and PFC ^2,6,30-32^. Thus, we assessed extracellular dopamine levels in these two terminal regions before and after activation of VTA neurons. We found that electrical and optogenetic stimulation of VTA neurons, using patterns of activation that resemble the phasic bursting activity of dopamine neurons during reward-guided behavioral tasks, results in increased extracellular levels of dopamine in the NAc and PFC that persist for many minutes after stimulation is terminated. This sustained post-stimulus elevation of dopamine is blocked by tetrodotoxin (TTX), suggesting that it is not due to dopamine spillover from the initial stimulation; rather, it results from reduced dopamine transporter (DAT)-mediated clearance of dopamine released spontaneously during the post-stimulation period.

## Results

### Electrical stimulation of VTA produces sustained dopamine release in NAc and PFC

Microdialysis was used to measure the extracellular concentration of dopamine ([DA]_∘_) in NAc and medial prefrontal cortex (mPFC) of freely moving rats during the active phase of their sleep-wake cycle in a home-cage environment. Microdialysis has a limit of detection in the femtomolar range ^33^, which is 2-3 orders of magnitude better than the detection limit of FSCV ^34^. The improved limit of detection afforded by microdialysis permits measurement of resting baseline [DA]_∘_ in behaving animals, including in regions with sparse dopamine innervation such as PFC ^20^. We were, therefore, able to detect resting [DA]_∘_ in NAc and mPFC before stimulation of VTA neurons and follow [DA]_∘_ until its return to a stable pre-stimulation level.

VTA was electrically stimulated after stable [DA]_∘_ was measured in NAc and mPFC. The electrical stimulation protocol (1 ms pulses delivered at 100 Hz for 200 ms, with an interburst interval of 500 ms) mimicked the bursting firing pattern observed in VTA dopamine neurons when rats are engaged in relevant behaviors. For example, in response to reward during a reward-guided operant task, VTA dopamine neurons (identified based on response to apomorphine) fire on average 1.1 bursts per second (interburst interval =∼ 900 ms) with a typical intraburst frequency of ∼30 Hz, but they can fire as fast as 100 Hz and higher ^35^. Hence, our protocol utilized interburst and intraburst frequencies that are within the range of reported frequencies. Furthermore, a similar 100 Hz stimulation protocol has been shown to evoke phasic (measured with FSCV) dopamine release in PFC with similar temporal dynamics and greater maximum dopamine concentration compared to a 20 Hz stimulation protocol ^12^, suggesting that the 100 Hz stimulation protocol is physiologically relevant. The stimulation duration was 5 to 20 min to model the typical duration of reward-driven behavioral paradigms ^20,22^. We found that electrical stimulation of VTA increased [DA]_∘_ in NAc and mPFC, and these elevations were sustained from several minutes to more than an hour post stimulation (Fig. 1, 2, and Supplementary Fig. 1, 2). A second stimulation given 140 min later produced a similar pattern of [DA]_∘_ increase (Fig. 1,2), suggesting that the initial stimulation had not provoked a pathological condition, such as sustained depolarization block of dopamine neurons ^36^. In addition, the magnitude and duration of dopamine response in both regions were similar to previously reported microdialysis measures in animals performing a reward-guided reversal learning task ^20^. These observations indicate that the pattern of stimulation used here is relevant to dopamine cell firing and dopamine release during reward-directed learning and cognitive behaviors. For subsequent studies, we focused on characterizing the impact of 20 min VTA stimulation on [DA]_∘_.

**Figure 1.**
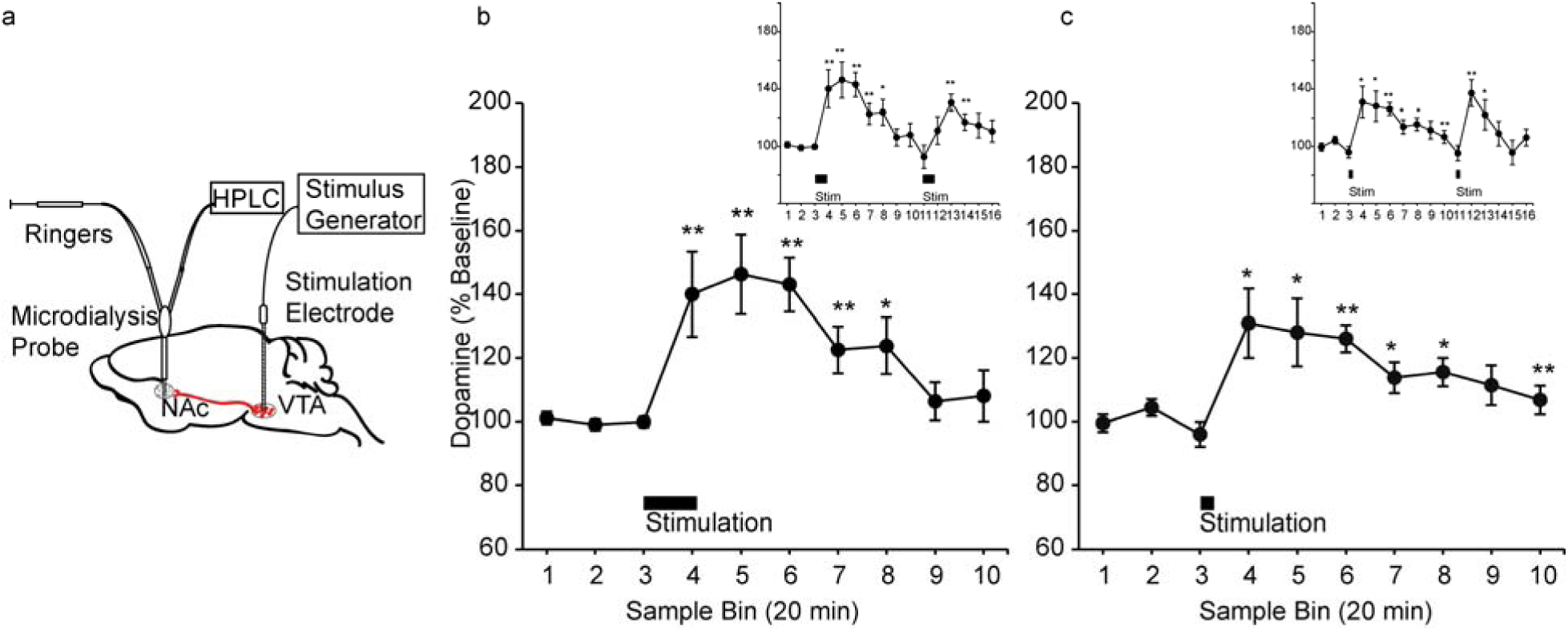
Electrical stimulation of VTA produces sustained dopamine release in NAc. (**a**) Schematic illustrating simultaneous measurement of [DA]_∘_ in NAc via a microdialysis probe and electrical stimulation of VTA via a stimulation electrode. (**b**) VTA was stimulated using a protocol consisting of bursts of 1 ms pulses delivered at 100 Hz for 200 ms, with an interburst interval of 500 ms and an amplitude of 60 μA, for 20 min (n = 10 sessions from 10 rats). VTA activation produced a sustained increase in [DA]_∘_ in NAc [F(15, 135) = 6.13, p = 0.00]. Post hoc comparisons revealed a significant difference between baseline (sample 3) and sample 4 (p = 0.01), sample 5 (p = 0.01), sample 6 (p = 0.00), sample 7 (p = 0.01), and sample 8 (p = 0.03). The figure inset illustrates dopamine release following a second delivery of 100 Hz stimulation. The second stimulation also significantly increased [DA]_∘_ in NAc in samples 13 (p = 0.00) and 14 (p = 0.01). (**c**) VTA activation using a stimulus protocol consisting of bursts of 1 ms pulses delivered at 100 Hz for 200 ms, with an interburst interval of 500 ms and an amplitude of 60 μA, for 5 min (n = 6 sessions from 6 rats) was associated with a sustained increase in [DA]_∘_ in NAc [F(15, 75) = 4.93, p = 0.00]. Post hoc comparisons revealed a significant difference between baseline (sample 3) and sample 4 (p = 0.03), sample 5 (p = 0.02), sample 6 (p = 0.00), sample 7 (p = 0.04), sample 8 (p = 0.02), and sample 10 (p = 0.01). The figure inset illustrates dopamine release following a second delivery of 100 Hz stimulation. The second stimulation resulted in an immediate elevation of [DA]_∘_ in NAc in sample 12 (p = 0.01) and sample 13 (p = 0.03). Data are represented as mean ± SEM. * p ≤ 0.05; ** p ≤ 0.01 (See also Supplementary Fig. 1)

**Figure 2.**
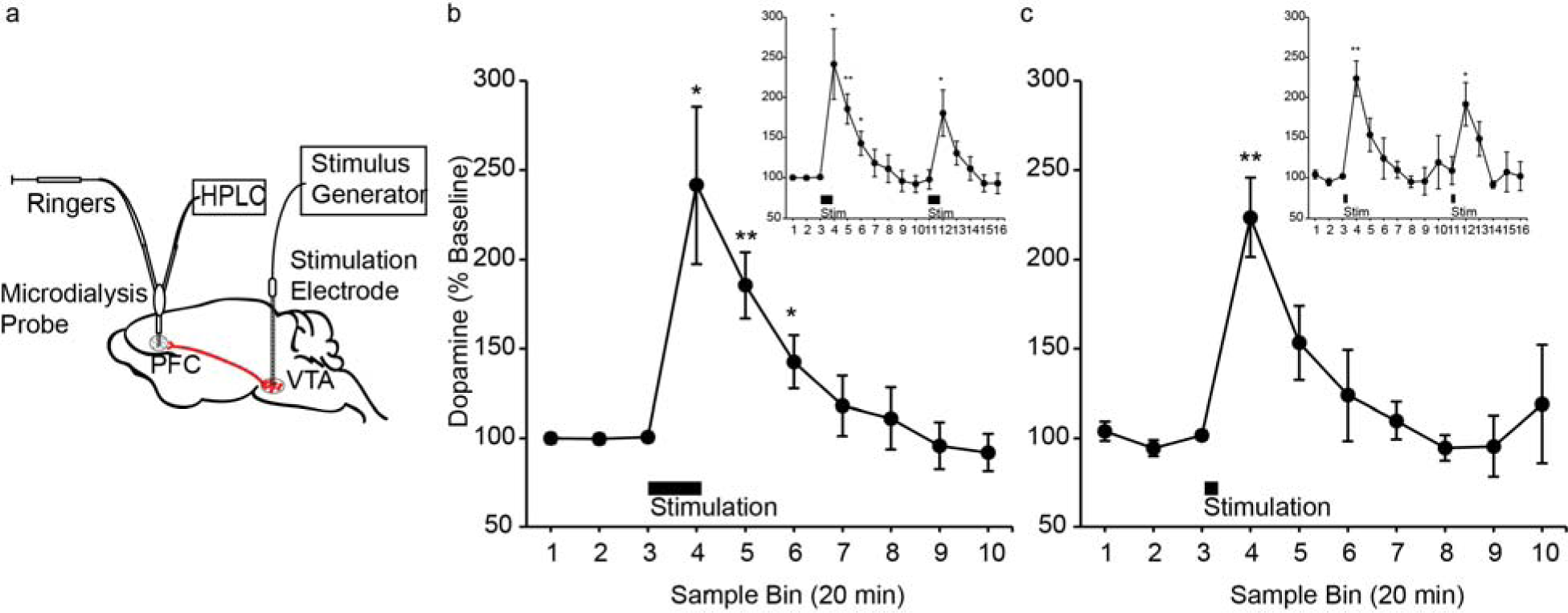
Electrical stimulation of VTA produces sustained dopamine release in PFC. (**a**) Schematic illustrating simultaneous measurement of [DA]_∘_ in PFC via a microdialysis probe and electrical stimulation of VTA via a stimulation electrode. (**b**) VTA stimulation using a protocol consisting of bursts of 1 ms pulses delivered at 100 Hz for 200 ms, with an interburst interval of 500 ms and an amplitude of 60 μΑ, for 20 min (n = 7 sessions from 7 rats) elevated [DA]_∘_ in PFC [F(15, 90) = 8.10, p = 0.00]. Post hoc comparisons revealed a significant difference between baseline (sample 3) and sample 4 (p = 0.02), sample 5 (p = 0.01) and sample 6 (p = 0.04). The inset illustrates dopamine release following a second delivery of 100 Hz stimulation, which significantly increased [DA]_∘_ in PFC in sample 12 when compared to baseline (p = 0.04). (**C**) VTA activation using a stimulus protocol consisting of bursts of 1 ms pulses delivered at 100 Hz for 200 ms, with an interburst interval of 500 ms and an amplitude of 60 μA, for 5 min (n = 6 sessions from 6 rats) was associated with sustained elevations in [DA]_∘_ in PFC [F(15, 75) = 5.15, p = 0.00]. Post hoc comparisons against baseline indicated a significant increase in [DA]_∘_ in sample 4 (p = 0.00). The inset illustrates dopamine release following a second delivery of 100 Hz stimulation. Post hoc comparisons revealed that [DA]_∘_ in PFC in sample 12 was significantly increased compared to baseline (p = 0.02). Data are represented as mean ± SEM. * p ≤ 0.05; ** p ≤ 0.01 (See also Supplementary Fig. 2)

### Optogenetic stimulation of VTA dopamine neurons produces sustained dopamine release in NAc

In addition to dopamine neurons, VTA contains a significant portion of GABA neurons ^37^, which could have been activated by VTA electrical stimulation. While it is unlikely that co-activation of GABA neurons in the VTA produces terminal release of dopamine, we nonetheless compared dopamine responses in NAc during and after electrical stimulation of VTA neurons to dopamine responses in NAc during and after selective optogenetic stimulation of VTA dopamine neurons in a tyrosine hydroxylase (*Th*)*::Cre* rat line ^38^. Using recombinant adeno-associated viral vectors, we expressed Cre-inducible channelrhodopsin-2 (ChR2) proteins in VTA of *Th::Cre* rats. To stimulate VTA dopamine neurons, we delivered blue laser (∼473 nm) into VTA using one of two protocols: a) 1 ms pulses delivered at 100 Hz for 200 ms, with an interburst interval of 500 ms, for 20 min (identical to the electrical stimulation protocol) or b) 5 ms pulses delivered at 20 Hz for 5 s, with an interburst interval of 10 s, for 20 min. Similar to electrical stimulation of VTA, optogenetic stimulation of VTA dopamine neurons elicited prolonged increases in [DA]_∘_ in NAc that persisted, on average, for 20 min after the cessation of stimulation (Fig. 3 and Supplementary Fig. 3).

**Figure 3.**
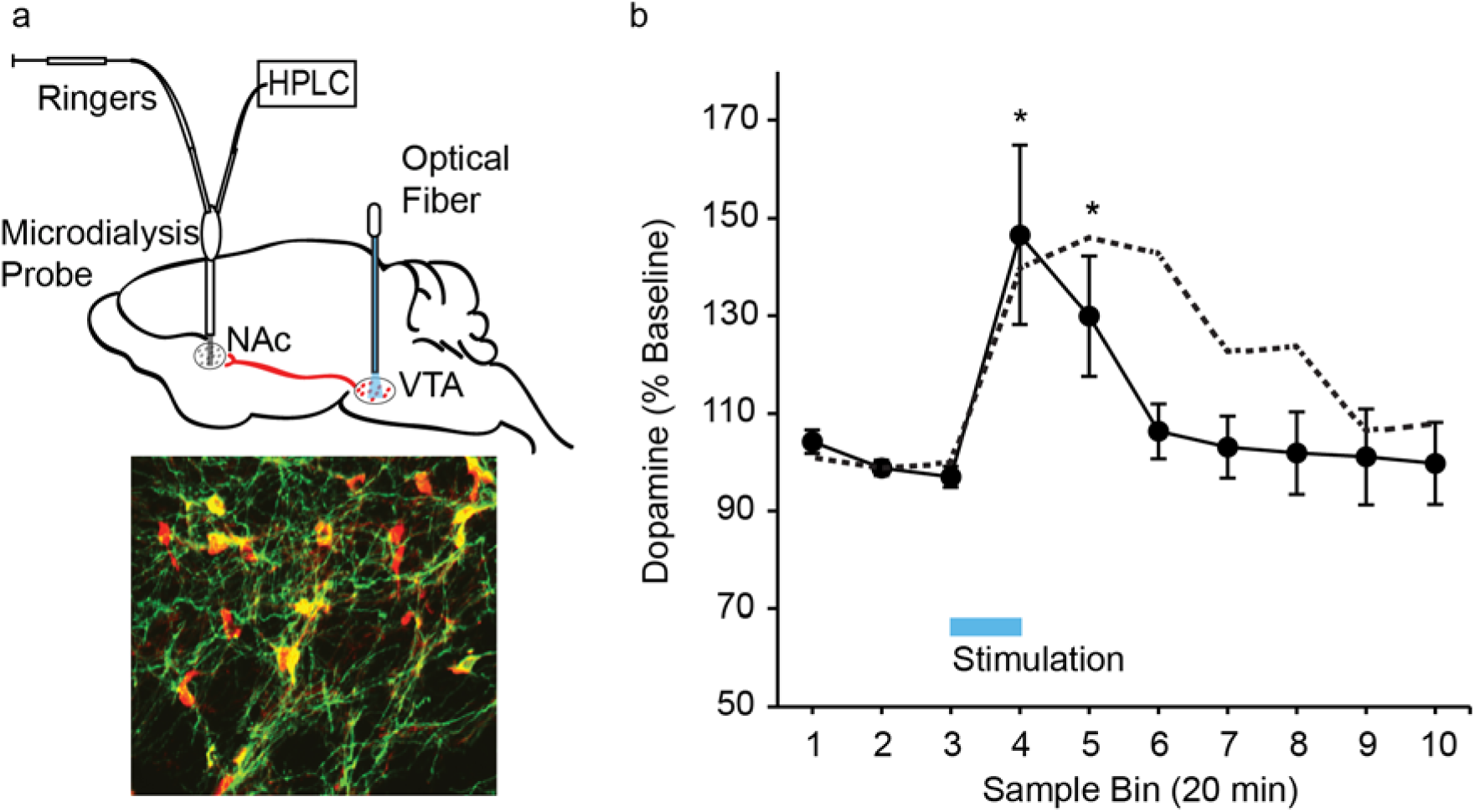
Optogenetic stimulation of VTA dopamine neurons produces sustained dopamine release in NAc. (**a**) (Top) Schematic illustrating simultaneous measurement of [DA]_∘_ via a microdialysis probe in NAc and blue laser illumination of ChR2-expressing VTA dopamine neurons via an optical fiber. (Bottom) TH (red) and ChR2-eYFP (green) immunofluorescence in VTA illustrates colocalization within VTA dopamine neurons (yellow) in a representative *Th::Cre* rat. (**b**) VTA dopamine neurons were stimulated using either a protocol consisting of bursts of 1 ms pulses delivered at 100 Hz for 200 ms, with an interburst interval of 500 ms, for 20 min or a protocol consisting of bursts of 5 ms pulses delivered at 20 Hz for 5 s, with an interburst interval of 10 s, for 20 min (laser power = 5-7 mW) (n = 11 sessions from 5 rats). Optogenetic VTA dopamine stimulation was associated with a sustained increase in [DA]_∘_ in NAc [F(9, 90) = 4.47, p = 0.00]. Post hoc comparisons revealed a significant difference between baseline (sample 3) and sample 4 (p = 0.02) and sample 5 (p = 0.02). The dotted line represents electrical stimulation-induced elevations in NAc [DA]_∘_ (depicted also in Fig. 1b). Data are represented as mean ± SEM. * p ≤ 0.05 (See also Supplementary Fig. 3)

### Post-stimulation sustained [DA]_∘_ increase in NAc and PFC depends on active release

Next, we assessed whether the observed sustained elevation of post-stimulation [DA]_∘_ in mPFC and NAc resulted from continued active release of dopamine or simply reflected remaining dopamine that was “leftover” after the large burst-induced release. Previous studies have shown that stimulation of dopamine neurons in a burst pattern produces greater release of dopamine than tonic stimulation, even when the number of impulses per second and the total number of impulses are held constant ^39,40^. Thus, burst firing of dopamine neurons can simply produce a large surge of [DA]_∘_ that clears away slowly. We found that blockade of voltage gated sodium channels via local application of TTX to either NAc or mPFC immediately after VTA stimulation drastically reduced [DA]_∘_ (Fig. 4 and Supplementary Fig. 4). In fact, [DA]_∘_ in NAc after TTX application approached the lower limit of detection in post-stimulation samples (mean of sample 6: 20 ± 7% and sample 7: 16 ± 7%) whereas stimulation in the absence of TTX continued to produce significant elevations in [DA]_∘_ (mean of sample 6: 143 ± 8% and sample 7: 123 ± 7%). Similarly, in PFC, mean [DA]_∘_ after TTX treatment fell to 19 ± 5% of baseline by post-stimulation sample 6 while stimulation alone still elicited a significant [DA]_∘_ increase at that time point (mean: 143 ± 15%). While some of the reduction in post-TTX [DA]_∘_ can be explained by TTX’s effect on background (baseline) extracellular dopamine, the fact that extracellular NAc and PFC dopamine levels dropped close to the detection limit suggests that TTX also prevented most of the sustained increase in [DA]_∘_ elicited by VTA stimulation. These results indicate that the sustained post-burst elevation of [DA]_∘_ may be supported by active (impulse-flow-dependent) exocytotic release of dopamine.

**Figure 4.**
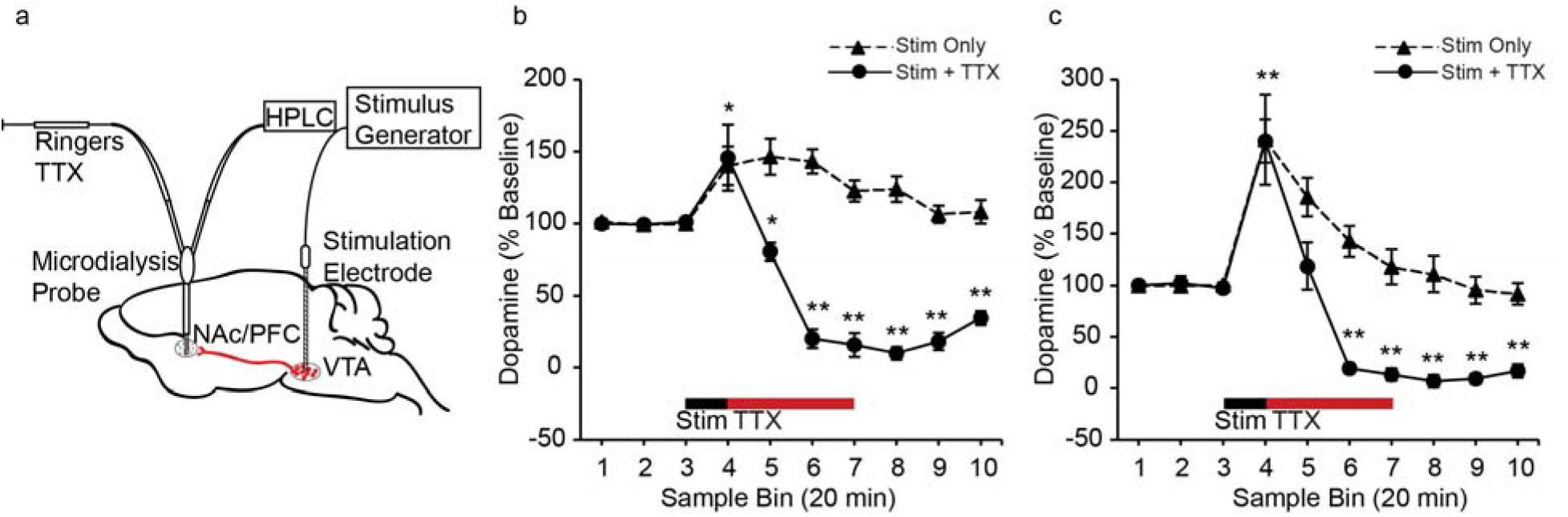
TTX blocks sustained dopamine release in NAc and PFC. (**a**) Schematic of infusion of TTX via microdialysis probes into NAc or PFC after electrical stimulation of VTA. (**b**) TTX infusion in NAc after cessation of VTA electrical stimulation blocked the post-stimulation increase in [DA]_∘_ in NAc. The solid line indicates data from sessions where TTX infusion in NAc followed electrical stimulation of VTA (stim + TTX, n = 7 sessions from 7 rats) while the dotted line represents data from rats that received electrical stimulation of VTA only (stim only, presented also in Fig.1b). Stim only and stim +TTX data were analyzed after applying log transformations (see Methods). Electrical stimulation of VTA alone produced sustained increases in [DA]_∘_ (results after data transformation were identical to the results in Fig. 1b; one way repeated measures ANOVA: F(9,81) = 5.93, p = 0.00; post-hoc tests against baseline: sample 4 (p = 0.01), sample 5 (p = 0.01), sample 6 (p = 0.00), sample 7 (p = 0.01), and sample 8 (p = 0.04)). TTX treatment modulated stimulation-induced increases in [DA]_∘_ in NAc (one way repeated measures ANOVA, F(9, 54) = 24.46, p = 0.00). Post hoc comparisons against baseline (sample 3) indicated that VTA stimulation before TTX infusion in NAc significantly increased NAc [DA]_∘_ in sample 4 (p = 0.04). Following TTX treatment, [DA]_∘_ decreased below baseline in sample 5 (p = 0.02), sample 6 (p = 0.01), sample 7 (p = 0.00), sample 8 (p = 0.00), sample 9 (p = 0.00) and sample 10 (p = 0.00). A two-way repeated measures ANOVA further demonstrated a significant interaction between sample bin and condition (stim + TTX and stim only) [F(9, 135) = 33.77, p = 0.00]. (**c**) TTX infusion in PFC after cessation of VTA electrical stimulation blocked the post-stimulation increase in [DA]_∘_ in PFC. As in panel B, the solid line indicates stim + TTX condition (n = 6 sessions from 6 rats) and the dotted line represents electrical stimulation of VTA only (stim only, presented also in Fig. 2b). TTX treatment after VTA electrical stimulation modulated sustained increases in [DA]_∘_ (one way repeated measures ANOVA: F(9,45) = 45.35, p = 0.00). Post-hoc comparisons against baseline (sample 3) indicated that while stimulation increased [DA]_∘_ in sample 4 (p = 0.00), [DA]_∘_ was no longer significantly different from baseline in sample 5 (p = 0.40) after TTX infusion in NAc. Further, [DA]_∘_ in PFC decreased below baseline in the remaining post-stimulation samples 5, 6, 7, 8, 9 and 10 (p = 0.00 for all comparisons). A two-way repeated measures ANOVA also demonstrated a significant interaction between sample bin and condition (stim + TTX and stim only) [F(9, 99) = 6.79, p = 0.00]. * p ≤ 0.05 and ** p ≤0.01 for comparison against baseline within the stim + TTX condition only. Data are represented as mean ± SEM. (See also Supplementary Fig. 4)

### Blockade of cannabinoid and nicotinic receptor activity does not impact post-stimulation increase in NAc [DA]_∘_

The outcome of the TTX experiment suggested that the burst-induced increase in [DA]_∘_ triggers secondary events that enhance active exocytotic dopamine release and/or reduce the clearance of dopamine actively released in the post-stimulation period. In the striatum, several secondary mechanisms that locally regulate release of dopamine have been identified ^41,42^. For example, an increase in synaptic dopamine release and subsequent dopamine receptor stimulation mobilizes the release of endocannabinoids ^43^ that could activate type-1 cannabinoid receptors (CB1Rs) on GABA or other neurons that may modulate dopamine release ^41^. We found that inhibition of CB1R activity via systemic injection of rimonabant did not change spontaneous dopamine levels or the pattern of stimulation-induced increase in [DA]_∘_ in NAc (Supplementary Fig. 5).

Another potential mechanism is secondary activation of cholinergic neurons and subsequent stimulation of presynaptic nicotinic acetylcholine receptors (nAChRs) on dopamine axons, evoking dopamine release that is independent of dopamine somal activity ^44,45^. To test this, the effect of local inhibition of nAChRs by mecamylamine on NAc [DA]_∘_ was examined. The results (Supplementary Fig. 6a,b) of the experiment did not provide support for this mechanism. Mecamylamine treatment in combination with VTA stimulation also elicited a large variability in post-stimulation samples, which could be due to the impact of mecamylamine treatment on basal [DA]_∘_ (Supplementary Fig. 6c,d).

### Sustained increase in [DA]_∘_ post-stimulation results from reduced DAT-mediated clearance of dopamine

Next, we considered mechanisms that are involved in the clearance of dopamine from the extracellular space. Previously, it has been shown that amphetamine, which acts as a substrate of DAT and causes a large surge of extracellular dopamine, induces transient internalization of DATs ^46^. Although microscopy studies have not been able to confirm this internalization ^47^, other recent work indicates that the process is mediated, in part, by activation of Rho-GTPases, since selective inhibition of Rho or stimulation of protein kinase A (PKA)-coupled receptors such as D1/5 dopamine receptors prevents amphetamine-induced DAT internalization ^48^. We hypothesized that a surge in synaptic dopamine release caused by burst activation of cell bodies also results in DAT internalization. Such a mechanism would then lead to an increase in [DA]_∘_ even after cessation of burst activation of dopamine neurons by reducing the rate of clearance of newly released dopamine. We found that dopamine at a high concentration (10 μΜ) decreased cell surface expression of DATs in rat midbrain slices (Fig. 5a). The same concentration of dopamine increased the amount of activated Rho GTPase in midbrain slices (Fig. 5b). We then tested the effects of inactivating Rho on dopamine-mediated DAT internalization. To inactivate Rho, we stimulated D1/5 receptors in midbrain slices with SKF 38393. D1/5 receptors are G-protein coupled receptors (GPCRs) present on dopamine neurons, and stimulation of these GPCRs increases adenylyl cyclase activity and cyclic adenosine monophosphate production, which activates PKA ^49^. Activated PKA phosphorylates and thus inactivates Rho ^50^. We found that SKF 38393 blocked dopamine’s effects on Rho-activation and DAT internalization (Fig. 5a,b).

**Figure 5.**
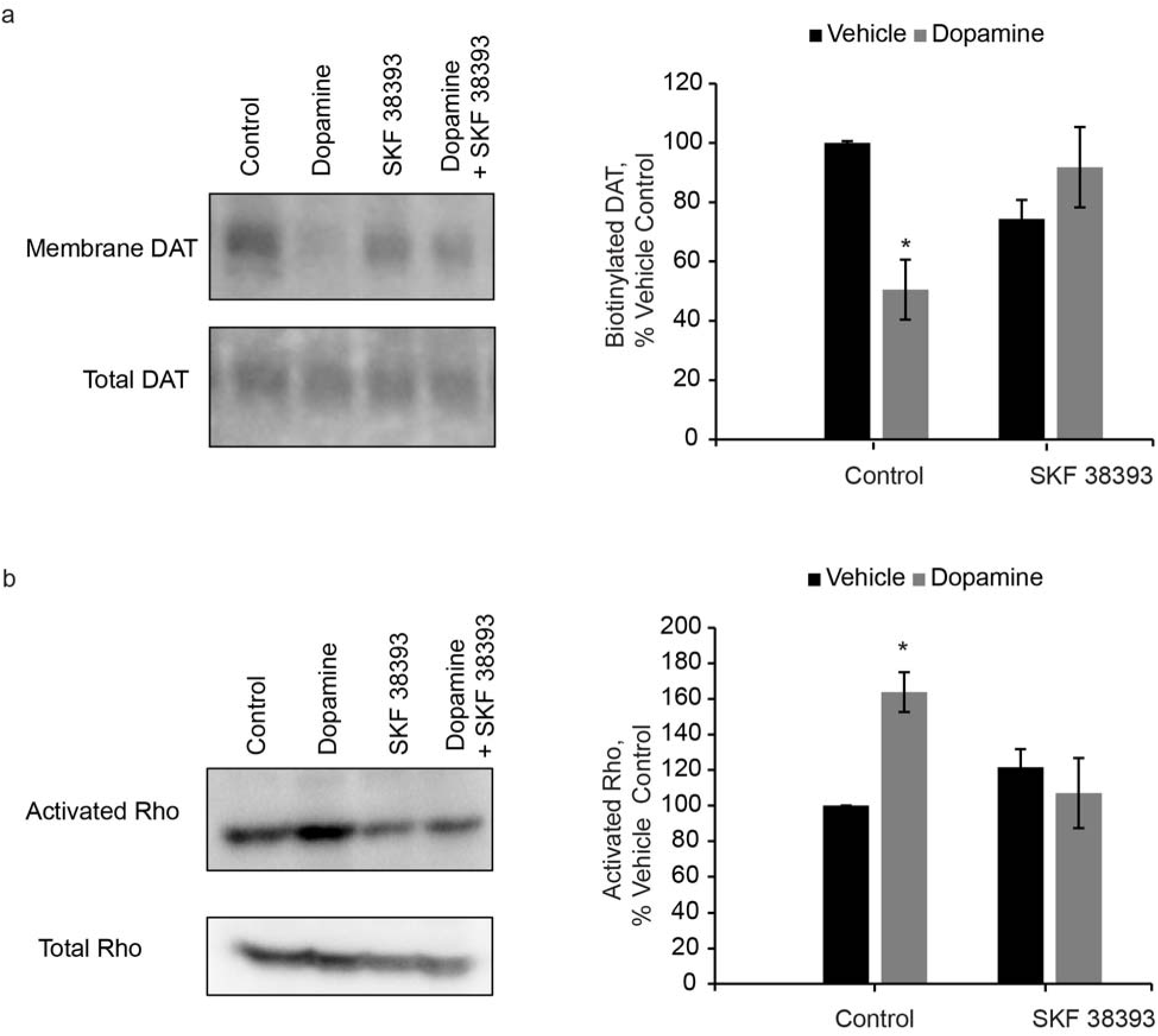
Stimulation of D1/D5 receptor blocks dopamine-induced decrease in cell surface expression of DAT and increase in Rho activation. Rat midbrain slices were treated with dopamine (10 μM), SKF 38393 (100 nM), or dopamine and SKF 38393. (**a**) Cell surface proteins were biotinylated, isolated and probed for cell surface expression. The effect of dopamine on membrane-localized DAT depended on SKF treatment [F(1, 8) = 13.72, p = 0.01, dopamine X SKF 38393 interaction, n = 3 slices for all groups]. Post hoc comparisons against control slices treated with vehicle revealed that membrane-localized DAT was significantly decreased in response to dopamine (p = 0.04), but this effect was blocked by SKF38393 (p = 2.01). SKF38393 treatment alone did not significantly increase dopamine levels but resulted in a non-significant trend towards a decrease in membrane-localized DAT (p = 0.06). (**b**) Tissue lysates were then assessed for activated Rho. A two-way ANOVA revealed a significant interaction between dopamine and SKF 38393 treatments [F(1, 12) = 10.00, p = 0.01] (n = 4 slices all groups). Post hoc comparisons against control slices treated with vehicle indicated that dopamine increased the amount of activated Rho GTPase in midbrain slices (p = 0.01), and this effect was blocked by SKF38393 (p = 0.75). SKF 38393 alone did not alter the amount of activated Rho-GTPase (p = 0.12). Data are represented as mean ± SEM. * p ≤ 0.05.

To establish the relevance of a Rho-mediated DAT internalization mechanism in freely moving animals, we injected rats with SKF 38393 (3 mg/kg, i.p.) prior to VTA electrical stimulation. This treatment prevented any VTA stimulation-induced increase in [DA]_∘_ in NAc (Fig. 6a). As this effect could result from the activation of D1/5 receptors elsewhere in the brain upon systemic injection of SKF 38393, we conducted a separate experiment where we infused SKF 38393 in NAc to assess the effect of local D1/5 receptor blockade on VTA stimulation-elicited increase in NAc [DA]_∘_. We found that local SKF 38393 infusion also attenuated sustained increases in dopamine levels in post stimulation samples (Fig. 6b). Hence, our data suggest that blockade of DAT internalization by SKF 38393 interrupts a sustained post-stimulation increase in [DA]_∘_. We found that SKF38393 treatment alone (i.p. and local infusion in NAc) also decreased [DA]_∘_ in NAc (Supplementary Fig. 7). However, there was no difference in [DA]_∘_ between SKF only and SKF + stimulation conditions in post-stimulation samples, suggesting that SKF 38393 prevented stimulation-induced sustained increases in [DA]_∘_ over a new drug-induced [DA]_∘_ baseline. Further, these results indicate that SKF 38393 might reduce DAT internalization in both baseline and stimulation conditions in freely moving animals.

**Figure 6.**
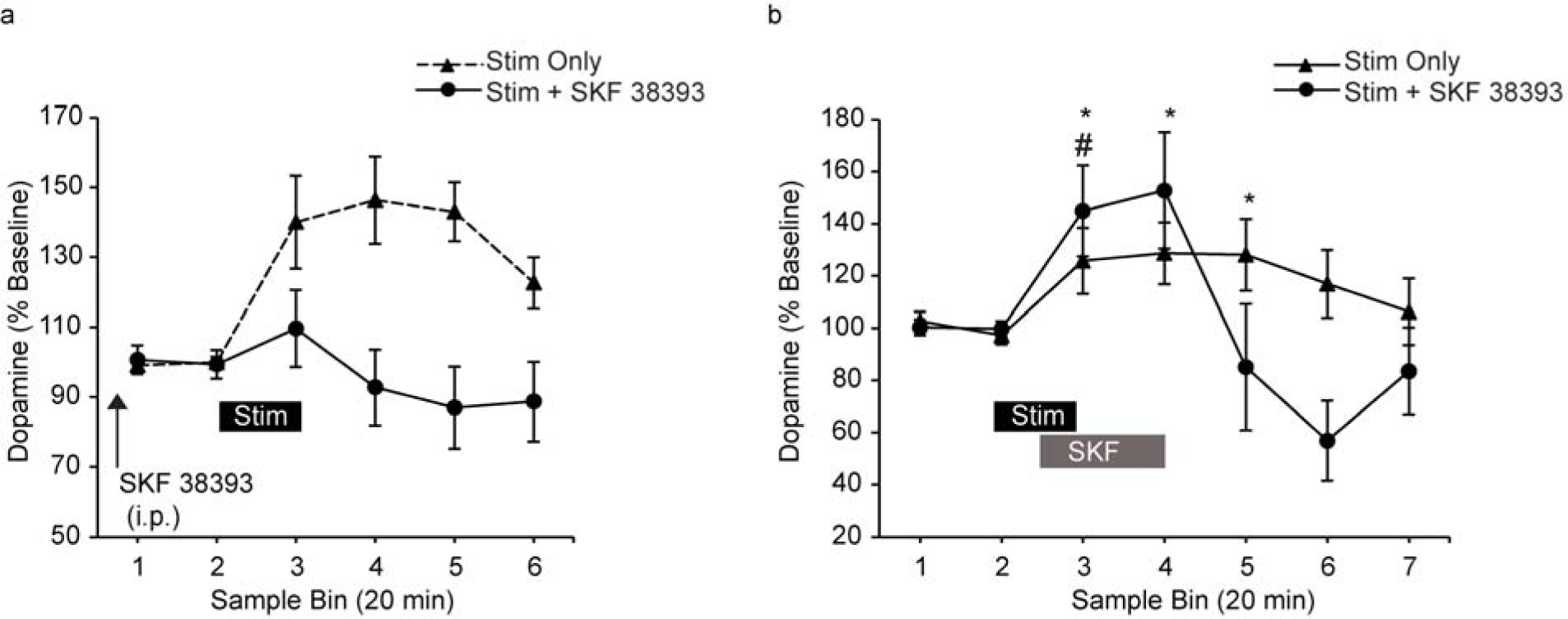
D1/D5 receptor activation attenuates sustained dopamine release in NAc. (**a**) Pre-treatment with SKF 38393 (i.p, 3 mg/kg) attenuated VTA stimulation-induced elevations in [DA]_∘_ in NAc. The solid line indicates data from sessions with SKF 38393 injection and electrical stimulation of VTA (stim + SKF 38393, n = 5 sessions from 5 rats) and the dotted line represents data from sessions with stimulation of VTA only (stim only, also presented in Fig. 1b). A one-way repeated measures ANOVA revealed that SKF 38393 prevented any change in [DA]_∘_ upon VTA stimulation (F(5, 20) = 1.12, p = 0.38). A two-way repeated measures ANOVA further established a significant difference between SKF 38393 + stim and stim only conditions (sample bin X condition: F(5, 65) = 4.19, p = 0.00). (**b**) Local infusion of SKF 38393 (10 μM) in NAc 10 min after the start of electrical stimulation of VTA attenuated post-stimulation increase in [DA]_∘_ in NAc. Filled triangles represent data from sessions with electrical stimulation of VTA only (stim only, n = 5 sessions from 5 rats) and filled circles represent data from sessions with SKF 38393 infusion in NAc and electrical stimulation of VTA (stim + SKF 38393, n = 5 sessions from 5 rats). For these sessions, VTA was stimulated using a protocol consisting of bursts of 1 ms pulses delivered at 100 Hz for 200 ms, with an interburst interval of 500 ms and an amplitude of 5 μA (compared to 60 μA used in previous experiments, see Methods), for 20 min. Electrical stimulation of VTA increased [DA]_∘_ in NAc (one way repeated measures ANOVA, F(6, 24) = 3.53, p = 0.01). Post hoc comparisons against baseline (sample 2) demonstrated significant elevations of [DA]_∘_ in sample 3 (p = 0.05), sample 4 (p = 0.03), and sample 5 (p = 0.05). Electrical stimulation of VTA in combination with SKF 38393 treatment also resulted in a significant modulation of NAc [DA]_∘_ (one way repeated measures ANOVA, F(6,24) = 6.29, p = 0.00). Post hoc comparisons against baseline (sample 2) demonstrated that even in the presence of SKF 38393 in NAc, electrical stimulation of VTA produced a significant elevation in NAc [DA]_∘_ in stimulation sample 3 (p = 0.04) and resulted in a trend towards a significant [DA]_∘_ increase in post-stimulation sample 4 (p = 0.08). However, SKF 38393 treatment prevented a sustained stimulation-induced [DA]_∘_ increase in sample 5 (p = 0.58) and resulted in a trend towards a decrease in [DA]_∘_ below baseline in sample 6 (p = 0.06). * and # indicate p ≤ 0.05 for comparisons against baseline within stim only and stim + SKF 38393 conditions respectively. Further, a two way repeated measures ANOVA confirmed that the local blockade of D1/5 receptors in NAc by SKF38393 changed the pattern of [DA]_∘_ increase elicited by electrical stimulation of VTA (sample X condition: F(6, 48) = 4.23, p = 0.00). Data are represented as mean ± SEM. (See also Supplementary Fig. 7,8)

**Figure 7.**
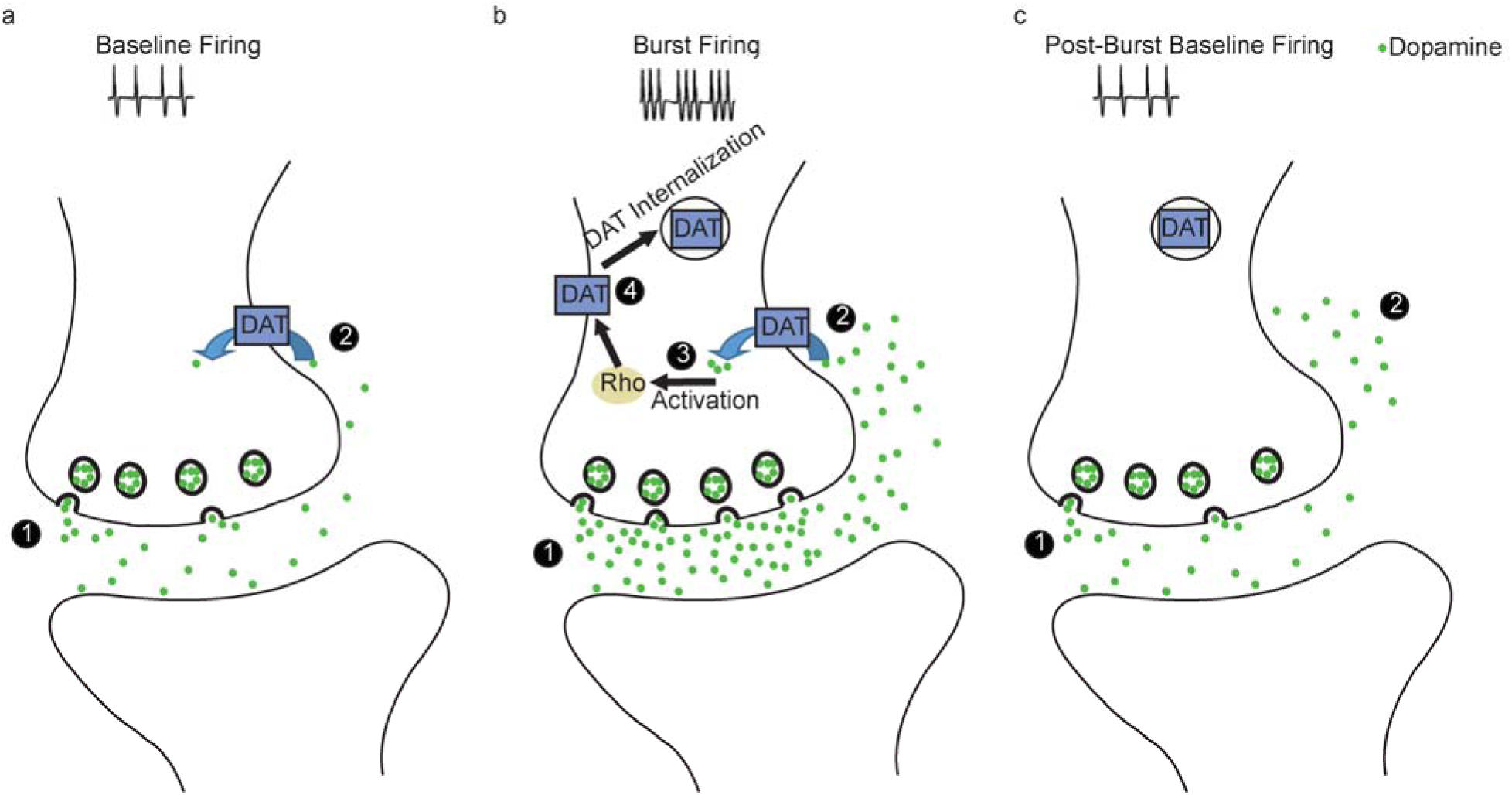
Mechanism for sustained increase in extracellular dopamine following burst activation of dopamine neurons. (**a**) 1) Baseline spontaneous firing of dopamine neurons causes the release of low levels of dopamine from the terminals. 2) Dopamine diffuses into the extrasynaptic space where it is taken up by DATs, resulting in a low level of extracellular dopamine. **(b)** 1) Dopamine neurons fire in phasic bursts in response to behaviorally salient stimuli. Phasic bursting results in increased release and higher synaptic levels of dopamine. 2) A high concentration of extracellular dopamine enters the cell via DATs. 3) Intracellular dopamine activates Rho. 4) Rho activation mediates internalization of DATs. (**c**) 1) Upon cessation of phasic bursting, dopamine neurons return to baseline firing and release low levels of dopamine into the synapse. 2) Synaptic dopamine diffuses into the extrasynaptic space, where very few DATs are present following DAT internalization. As a result, extrasynaptic dopamine is cleared away slowly and dopamine accumulates in the extracellular space.

## Discussion

We found that phasic burst activity of VTA dopamine neurons in awake rats in a home-cage environment is followed by a sustained post-burst elevation of [DA]_∘_ in both NAc and PFC terminal regions that persisted for at least 20 min. This increase is not due to dopamine spillover from the initial stimulation but depends on continued active release of dopamine in terminal regions. We identified Rho-mediated internalization of DATs as a potential mechanism responsible for increased post-burst availability of actively released dopamine, based on evidence that experimental blockade of Rho activation prevented both DAT internalization and the sustained post-stimulation elevation of [DA]_∘_ in NAc. These results suggest that phasic surges of dopamine release produced by bursting of dopamine neurons result in DAT internalization that reduces the clearance of actively released dopamine in the post-burst period, causing a sustained post-burst elevation of tonic [DA]_∘_.

### Sustained post-burst increase in [DA]_∘_ is consistent with behavioral evidence and volume transmission

The sustained increase in [DA]_∘_ after periods of burst activity is functionally significant, because dopamine perisynaptic architecture in cortical and striatal regions is well suited to accommodate a slow and spatially distributed signal via volume transmission ^41,51^. A considerable density of dopamine receptors, both in the PFC and striatum, is localized extrasynaptically ^52-54^, and more than half of dopamine release sites in the striatum are reported to lack post-synaptic specializations ^55^. Hence, dopamine that slowly accumulates in the extracellular space on the order of minutes to hours could modulate the activity of many pre- and post-synaptic sites within a large area in the striatum and the PFC via volume transmission ^27^. This sustained and diffuse dopamine signal might maintain and stabilize/consolidate networks activated during learning and working memory ^56^. In addition, persistent dopamine signaling could lower the threshold for long term potentiation in all synapses for a certain period of time, allowing for adaptive modulation of memory encoding ^57^.

The mechanisms identified here might be especially critical for explaining dopamine’s role in memory consolidation during reinforcement-guided learning. Previous microdialysis studies in animals performing reward-driven learning or working memory tasks have demonstrated that [DA]_∘_ in PFC and striatal regions remains elevated for many minutes after completion of the training phase ^19-21^. In fact, the duration and magnitude of the post-training increase in [DA]_∘_ in these studies were similar to what we observed in the present study after electrical or optogenetic burst stimulation of dopamine neurons. The importance of this post-training dopamine signaling has been recognized ^24-26,58^, but the underlying mechanisms are unknown. Dopamine neurons fire phasically in response to salient stimuli during a behavioral task, but these phasic increases in activity return to baseline levels after hundreds of milliseconds ^78,59-62^. This raises the question of how elevated dopamine levels can be sustained post-training, when dopamine neurons are presumably firing at baseline levels. Our data obtained under electrical and optogenetic stimulation conditions suggest that repetitive, stimuli-evoked bursting of dopamine neurons during behavioral training may produce changes in DAT internalization processes, resulting in an extracellular accumulation of spontaneously released dopamine posttraining. This finding also reconciles rapid dopaminergic transmission during learning with slow and sustained dopamine response after learning.

Prolonged dopamine responses have also been implicated in sustained motivation ^4,63-66^. For example, sustained increases in extracellular dopamine that continue after the end of behavior are observed in the striatum when rats lever press for food or lick a spout for sucrose solution, and these prolonged dopamine elevations are modulated by shifts in motivational state ^23,66,67^. Furthermore, a computational model has indicated that a slow increase in striatal dopamine levels that might carry an averaged reward signal during a behavioral task predicts motivational state ^68^. Hence, sustained increases in [DA]_∘_, as observed in the current study, might be critical for plasticity mechanisms that include memory consolidation and adaptive learning as well as persistent functions such as working memory and motivation.

### Proposed mechanism of sustained dopamine release following burst activation

While phasic and sustained (or tonic) modes of dopamine neurotransmission are often hypothesized to be driven by distinct mechanisms and to affect different postsynaptic processes and behaviors ^27-29,69^, the present findings suggest that, in awake and active animals, these two modes of signaling may be complementary (Fig. 7). Ongoing activity of dopamine neurons in awake animals maintains impulse flow-dependent resting extracellular levels of dopamine in terminal regions such as the PFC and NAc (Fig.7a) ^70^. These levels are regulated by DAT, since the local application of DAT inhibitors profoundly enhances [DA]_∘_ ^71,72^. In response to salient events during behavioral tasks, dopamine neurons respond with brief bursts of spikes ^7,8,59,60,62^ that generate a large phasic surge of [DA]_∘_ ^39,40^. Our data indicate that large increases in [DA]_∘_ may cause DAT internalization via activation of Rho-GTPases. We propose that dopamine enters the cell via DATs to initiate this Rho-dependent intracellular signaling cascade (Fig. 7b), based on evidence for a similar mechanism of amphetamine-mediated internalization of DATs and EAAT3 glutamate transporters on dopamine neurons that requires the entry of amphetamine into the cytosol ^48,73,74^. The mechanism via which intracellular dopamine activates Rho is unclear. Candidate mechanisms include influx of calcium and subsequent calmodulin/CamkII activation or activation of intracellular GPCRs. These mechanisms have been proposed to explain how intracellular amphetamine activates Rho ^74^. DAT internalization has important consequences for the period following burst stimulation, which would be analogous to the immediate post-task period in learning and cognitive tasks. In the post-task period, dopamine neurons return to their spontaneously firing activation level, which generates a low level of synaptic dopamine. However, as a result of DAT internalization, dopamine is cleared at a much slower rate, leading to a sustained accumulation of dopamine in the extracellular space (Fig. 7c).

### Role of D1/5 receptor activation in dopamine-induced DAT internalization

Here we demonstrated the role of Rho-activation in dopamine-mediated DAT internalization by indirectly inhibiting Rho with D1/5 receptor activation via SKF 38393. Although other mechanisms that activate PKA could block Rho-mediated DAT internalization, we utilized this experimental approach because D1/5 receptors are physiologically relevant in dopamine-related behaviors. For example, mice lacking D5 receptors display a greater locomotor response to methamphetamine, a dopamine-activating stimulant, compared to wild-type mice ^75^, suggesting that D5 receptor activation might serve to curb excessive dopamine release and accumulation. Hence, we speculate that when there is a burst-induced release of a large amount of dopamine, activation of presynaptic D1/5 receptors by extracellular dopamine counteracts, to some extent, Rho-activation and DAT internalization mediated by dopamine transported into the cell. In other words, in dopamine terminals at baseline conditions, there might be a balance between processes that result in DAT internalization and those that interrupt DAT internalization. The former could be mediated by intracellular dopamine that activates Rho and the latter by extracellular dopamine that activates presynaptic D1/5 receptors. When a large surge of dopamine occurs during phasic activation of dopamine neurons, the balance might be shifted toward Rho-mediated DAT internalization. On the other hand, when most presynaptic D1/5 receptors are stimulated after systemic administration of SKF38393, activation of D1/5-coupled PKA signaling cascades could outcompete ongoing Rho activation. Consistent with this idea, our data indicate that SKF 38393 not only prevents burst-induced increases in [DA]_∘_ but also decreases baseline [DA]_∘_. Reduction of basal striatal dopamine levels by SKF 38393 has been reported by previous studies as well ^76,77^.

### Effects of DAT independent mechanisms on [DA]_∘_

It should be underscored that the DAT-mediated clearance mechanism we have identified here may be only one of the mechanisms that contribute to the sustained post-stimulation activation of [DA]_∘_. Additional mechanisms may include activation of feedback loops, such as activation of NAc medium spiny neurons that inhibit GABA neurons in VTA, resulting in the disinhibition of VTA dopamine neurons ^78^, and/or local signaling via non-dopamine receptors, including glutamate, GABA and nicotinic acetylcholine receptors ^41,42,44,45^. Although we examined the roles of CB1R activation and nAChR stimulation in regulating burst-induced activation of dopamine release and found no significant effects, we do not discount the potential involvement of other mechanisms in influencing post-burst dopamine accumulation.

### Concluding remarks

The present findings indicate that a critical function of burst activity of dopamine neurons may be to produce a persistent elevation of extracellular dopamine levels via an intracellular mechanism that promotes DAT internalization. The prolonged post-stimulation increase in dopamine levels may be essential for consolidation of memories during associative learning, maintenance of active networks in working memory, and sustenance of motivational states. Hence, identification of a mechanism that supports this sustained increase in extracellular dopamine has important implications for understanding learning and memory processes in the brain and also may provide insight into dysfunction of dopamine systems in disorders such as addiction and schizophrenia.

## Experimental Procedures

### Animals

Male Sprague-Dawley rats were used for electrical stimulation experiments while optical stimulation experiments utilized male Long-Evans *Th::Cre* rats (gifted by Dr. Karl Deisseroth) that express Cre recombinase in TH positive neurons. All rats were housed in a 12 hour reverse light/dark cycle with lights on at 7 pm. Experiments were conducted according to the ethical guidelines of the Institutional Animal Care and Use Committee at the University of Pittsburgh and the Animal Care and Use Committee at the NIH.

### Surgery

All animals were anesthetized with isoflurane during surgical implantation. Rats used in most of the electrical stimulation experiments were unilaterally implanted with homemade microdialysis probes in the PFC (AP = 3.4 mm from Bregma, ML = 0.7 mm, DV = 5.2 mm from skull, and probe membrane length = 3 mm) and NAc (AP = 1.7 mm from Bregma, ML = 1.1 mm, and DV = 8.4 mm from skull, and probe membrane length = 2-2.5 mm). Bipolar stimulating electrodes were also implanted in the VTA (AP = -5.3 mm from Bregma, ML = 0.8 mm, and DV = 8.3 mm from skull). Rats in SKF 38393 experiments were unilaterally implanted with guide cannulae (CMA Microdialysis) in the NAc (AP = 1.7 mm from Bregma, ML = 1. 8 mm, and DV = 4-5 mm from brain surface) and bipolar stimulating electrodes in VTA (AP = -5.4 - 5.5 mm from Bregma, ML = 0.7 - 0.8 mm, and DV = 7.3 mm from brain surface or 8.3 mm from skull).

In optical stimulation experiments, *Th::Cre* rats were unilaterally injected with Cre-inducible recombinant AAV viral vector constructs containing the gene encoding ChR2 (AAV-Ef1α-DIO-ChR2-eYFP, University of North Carolina Vector Core) in four different locations of VTA (AP = -4.9 mm and -5.7 mm from Bregma, ML= 0.7 mm, and DV = -7.2 mm and -8.4 mm from brain surface). At each site, 1μl of the virus was injected at a rate of 0.1 μL□min^-1^ using a microsyringe (Hamilton Co.) and a pump (World Precision Instruments). Following sufficient time for viral expression (at least 4 weeks), rats were implanted with microdialysis guide cannulae (CMA Microdialysis) targeting NAc (AP = 1.7 mm from Bregma, ML = 1.8 mm, DV = 4 mm from brain surface) and optical fibers (200-μm core diameter, 0.22 NA, Doric Lenses) targeting VTA (AP = -5.3 mm from Bregma, ML = 0.7 mm, and DV = 7.5 mm from brain surface).

### Microdialysis Procedures

All microdialysis experiments were conducted in freely moving animals. After surgery, animals were allowed to recover for at least 24 hours prior to the collection of the dialysate samples. For animals implanted with chronic guide cannulae, animals were allowed to recover for at least one week before starting experiments, and probes (CMA Microdialysis) of 3 mm membrane length were inserted into the cannulae on the day of the experiment. Probes were perfused with Ringer’s solution (in mM: 37.0 NaCl, 0.7 KCl, 0.25MgCl_2_, and 3.0 CaCl_2_) at a flow rate of 2.0 μL/min during sample collection. Dialysate samples were collected every 20 min and immediately injected into an HPLC system for electrochemical detection of dopamine as described before ^79^. After a stable baseline was established, treatment was delivered: VTA electrical stimulation alone, VTA optical stimulation alone, VTA electrical stimulation + TTX (Sigma, 1 μM), mecamylamine (Sigma, 100 μM) alone, mecamylamine (100 μM) + VTA electrical stimulation, rimonabant (NIDA, i.p., 1 mg/kg) alone, rimonabant (i.p., 1mg/kg) + VTA electrical stimulation, SKF 38393 (Tocris Bioscience, i.p.,3 mg/kg) alone, SKF 38393 (i.p., 3 mg/kg) + VTA electrical stimulation, SKF 38393 (Tocris Bioscience, 10 μM) alone, and VTA electrical stimulation + SKF 38393 (10 μM). In the VTA electrical stimulation only experiments, VTA was electrically stimulated twice for 20 min each. The second stimulation was delivered 140 min after the end of first stimulation. In the optical stimulation only experiments, VTA dopamine neurons were optogenetically stimulated once for 20 min. In the TTX experiments, TTX was mixed in Ringer’s solution and perfused directly into PFC or NAc through the microdialysis probes for 60 min after the cessation of VTA electrical stimulation. In the mecamylamine experiment, mecamylamine was mixed in Ringer’s solution and perfused into NAc through the microdialysis probes for 40 min at the start of VTA electrical stimulation. A separate group of rats received perfusions of mecamylamine in the NAc for 40 min but without VTA electrical stimulation. In the rimonabant + VTA electrical stimulation experiment, VTA was electrically stimulated 20 min after an i.p. injection of rimonabant. A separate group of rats received an i.p. injection of rimonabant alone without VTA electrical stimulation.

Two types of SKF 38393 experiments were conducted. In the first experiment, SKF 38393 was administered via an i.p. injection either alone or in combination with VTA electrical stimulation that started 60 - 80 min after drug injection. In the second experiment, SKF 38393 was mixed in Ringer’s solution and perfused directly into NAc through the microdialysis probes. The latter experiment was conducted using a counterbalanced within-subjects design, where each rat was given at least one session of each treatment (VTA stimulation only, VTA stimulation + SKF 38393, or SKF 38393 only) with at least one week of washout between sessions. In the stimulation only session, VTA electrical stimulation started 3 min prior to the end of a stable baseline sample and continued for 20 min. Electrical stimulation was shifted in time compared to sample collection in order to remove time lag in the measurement of dopamine response to stimulation because of dead volume in the probe and dialysis setup. As we corrected for dead volume, the first sample collected after the end of electrical stimulation was considered the stimulation sample. In the VTA stimulation + SKF 38393 session, SKF 38393 perfusion started 10 min into the stimulation sample and continued for 30 min. In the SKF 38393 only session, SKF 38393 was perfused for 30 min and VTA stimulation was not delivered.

### Stimulation Parameters

In electrical stimulation experiments, the implanted bipolar electrode was connected to a stimulator (Grass Technologies). VTA was then stimulated using parameters that mimic burst firing of dopamine neurons: 1 ms pulses delivered at 100 Hz for 200 ms, with an interburst interval of 500 ms. Stimulation was delivered for 5 or 20 min at an amplitude of 6 μA or 60 μA. 60 μA current was used for all experiments except for stimulation sessions in the second SKF 38393 (perfusion in NAc) experiment. 60 μA elicited behavioral activation in the former experiments (data not shown). In the latter experiment, 6 μA current was sufficient to elicit VTA stimulation-induced behavioral activation, and larger currents appeared to be too strong based on rats’ behavior (data not shown). Histology showed that stimulation electrode placements in these rats were more dorsal in the VTA compared to the previous experiments, which could explain the difference in stimulation current threshold that would elicit behavioral responses.

In optical stimulation experiments, a patch cord (Doric Lenses) attached to a 473-nm blue laser diode (OEM Laser Systems), which was controlled by a Master-8 stimulator (A.M.P.I), was connected to the implanted optical fiber. Blue laser (∼473 nm) pulses were delivered using one of two burst protocols: a) 20 pulses (pulse width = 1 ms) at 100 Hz for 200 ms, with an interburst interval of 500 ms; and b) 100 pulses (pulse width = 5 ms) at 20 Hz for 5 s, with an interburst interval of 10 s. Photostimulation was delivered for 20 min with light output between 5 - 7 mW (measured with a continuous pulse) at the tip of the optical fiber.

### Histology and Immunohistochemistry

Animals were anesthetized with chloral hydrate (400 mg/kg) and transcardially perfused with 0.9% saline and either 8% buffered formalin or 4% paraformaldehyde (PFA). Placements of microdialysis probes, stimulating electrodes and optical fibers were verified by staining coronal brain slices with cresyl-violet. Data from animals with incorrect placements are not included in this report.

To verify the expression of ChR2-eYFP in dopamine neurons, brains were fixed in 4% PFA overnight at 4°C and then transferred to a 20% sucrose solution for cryoprotection. Brains were frozen and sectioned at 35 μm. VTA coronal slices were then treated with 1% sodium borohydride in 0.1 M sodium phosphate buffer, rinsed with the buffer, and incubated in primary and secondary antibody solutions that contained 10% Triton-X and normal donkey serum dissolved in phosphate buffer. Dopamine cell bodies were labeled with mouse anti-tyrosine hydroxylase antibody (1:2000; EMD Millipore, #MAB318) followed by Cy3-tagged donkey antimouse antibody (1:500; Jackson ImmunoResearch). EYFP positive cells were identified using chicken anti-EGFP antibody (1:1000; Abcam, # ab13970) followed by Alexa Fluor 488-tagged donkey anti-chicken secondary antibody (1:500, Jackson ImmunoResearch). Expression of ChR2-eYFP in VTA dopamine neurons was confirmed by using a fluorescence microscope.

### Biochemistry of Acute Brain Slices

One mm midbrain slices were made from adult male Long Evans rats and maintained in artificial cerebral spinal fluid (in mM, 126 NaCl, 3.5 KCl, 2 CaCl_2_, 1.3 MgCl_2_, 1.2 NaH2PO4, 25 NaHCO3, 10 D-dextrose, pH 7.4) bubbled with 95% O_2_ / CO_2_ 5% at 37°C. Biotinylation assays were performed as described previously ^74^. Briefly, slices were treated for 30 min with the indicated drugs (dopamine 10 μM, SKF 38393 100 nM or dopamine and SFK 38393) and then rapidly chilled to 4°C to minimize further trafficking events. Cell surface proteins were biotinylated for 20 min with the cell-impermeable biotin reagent, sulfo-NHS-SS-biotin (Thermo Fisher Scientific), which was then quenched with glycine, 100 mM. The tissue was then lysed in avidin binding buffer (in mM, 150 NaCl, 5 EDTA, and 50 Tris, pH 7.5, with 1% Triton X-100 and a protease inhibitor). Biotinylated proteins were isolated by NeutrAvidin beads and analyzed by western blot. For Rho activation assays, slices were treated with the indicated drugs (dopamine 10 μM, SKF 38393 100 nM or dopamine and SFK 38393) for 5 min and rapidly chilled and lysed in rhotekin binding buffer (50 mM Tris Cl, pH 7.2, 1% Triton X-100, 500 mM NaCl, 10 mM MgCl_2_). GST conjugated to the Rho binding domain of rhotekin (RBP) was purified from *E. coli* and bound to glutathione-sepharose beads ^80,81^. Activated Rho proteins were captured in the tissue lysate by incubation with the GST-RBP beads overnight at 4°C. Active and bound Rho was eluted and analyzed by western blot. For both biotinylation and Rho activation assays, samples taken before purification were used to assess equal protein loading. All data were quantified with Image J.

## Data Analysis

In microdialysis experiments, dopamine concentration (fmol/μL) for each sample was expressed as a percentage of average baseline dopamine concentration. Two or three samples immediately before a treatment were considered to be baseline samples. If % baseline values fluctuated excessively within the same session, data from those sessions were not included.

All statistical analyses were conducted on % baseline values. The effect of a treatment (stimulation, drug alone or drug + stimulation) on PFC and NAc dopamine levels was assessed using a repeated measures ANOVA with sample bin as the within-subjects factor, and post hoc analyses were done with protected Fisher’s LSD tests to compare each sample bin with baseline. Two way repeated measures ANOVAs with sample bin as the within-subjects factor and treatment condition as the between subjects factor were also conducted to examine the difference between stimulation alone and stimulation + drug conditions. Before analyzing TTX + stimulation data from NAc only, data were log transformed (Y=log10(X+1)) because of robust violations of normality in multiple samples (based on Shapiro-Wilk test and examination of Q-Q plots). For comparison, stimulation only data from NAc were also reanalyzed after applying log transformations. To assess the difference between drug alone and stimulation + drug conditions, independent t-tests were utilized on a small subset of samples. The biochemistry data were analyzed using two way repeated measures ANOVAs followed by post-hoc t-tests. For all independent t-tests, equality of variances was assumed or not assumed based on the results of Levene’s test for equality of variances. All statistical values were rounded to decimal places and the null hypothesis was rejected if p ≤ 0.05.

## Data Availability

Any relevant data can be requested from the corresponding author.

## Author Contributions

A.M., A.D., S.L., B.M. and M.R. designed and performed the microdialysis experiments. S.A. and S.U. designed and performed the biochemistry experiments. L.R. performed the dual immunofluorescence localizations. A.M. and S.L. analyzed the data. B.M. and S.A. provided the financial support. S.L. and B.M. wrote the paper.

## Acknowledgments

This work was supported by the National Institutes of Health grant R01 (MH048404) to B.M and Intramural Research at the National Institute of Mental Health to S.A.

## Additional information

Supplementary Information accompanies this paper.

The authors declare no competing conflicts of interest.

